# The effect of acidic pH on the interaction and lytic activity of MP1 and its H-MP1 analog in anionic lipid membrane: a biophysical study by Molecular Dynamics and Spectroscopy

**DOI:** 10.1101/2020.12.11.404764

**Authors:** Ingrid Bernardes Santana Martins, Taisa Giordano Viegas, Bibiana Monson de Souza, Mário Sérgio Palma, João Ruggiero Neto, Alexandre Suman de Araujo

**Affiliations:** UNESP - São Paulo State University, IBILCE, Department of Physics, São José do Rio Preto, SP, Brazil; UNESP - São Paulo State Univeristy - Institute of Biosciences, Department of Basic and Applied Biology, Rio Claro, SP, Brazil

**Keywords:** Antimicrobial peptides, pH sensitive peptide, Molecular Dynamics Simulation, conformational analysis, lytic activity

## Abstract

Antimicrobial peptides (AMPs) are part of the innate immune system of many species and are compounds with potential application against the development of resistant bacterial strains promoted by conventional antibiotics. The AMPs are rich in cationic and hydrophobic residues and act directly on the lipidic phase of the cell membranes. The MP1 has a broad-spectrum bactericide activity in both Gram-negative and positive bacteria, not being hemolytic or cytotoxic. H-MP1 is a synthetic analog of MP1 with lysines replaced by histidines so that its net charge could be responsive to changes in solution pH. In the present work, we investigated the effect of the solution pH on the structural properties, in the adsorption and insertion, and on the lytic activity of these peptides in lipid bilayers mimicking the cell membrane of Gram-negative bacteria, using experimental and computational biophysical techniques. The results indicate that the lytic activity of H-MP1 is sensitive to pH, increasing to an acidic environment, matching that of MP1, which is not influenced by solution pH. Molecular Dynamic simulations indicated that the adsorption process of both peptides started by the interaction of the N-terminus with the bilayer, followed by the complete adsorption of the peptide laying parallel to the bilayer plane, inducing an increase in the peptide’s helical content enhancing peptides contact with the bilayer hydrophobic phase.

## 1 Introduction

Antimicrobial peptides (AMPs) are important alternatives to conventional antibiotics to face the emergence of resistant bacterial strains [1, 2]. These peptides are part of the innate immune response of many species from bacteria to man. By contrast to conventional antibiotics, AMPs act on the lipid-phase of plasma membrane without requiring membrane receptors [3, 4]. The development of resistance mechanisms by the bacteria can be hampered when the peptide acts on the lipid membrane [5]. AMPs are rich in non-polar and cationic residues that selectively enhance their preference for anionic lipid membranes, characteristic of prokaryotic organisms [6]. The tetradecapeptide Polybia–MP1, or simply MP1, extracted from the Brazilian wasp *Polybia paulista*, displays a potent broad-spectrum bactericide activity against both Gram-negative and Gram-positive bacteria [7]. Although it belongs to the family of mastoparan peptides it is not hemolytic neither cytotoxic and displays an inhibitory effect in cancer cell proliferation in cell culture[7].

An interesting characteristic of this peptide is the concomitant presence of two aspartic acids and four lysines distributed in such a way that each aspartic has lysines as its third or fourth neighbors in its sequence [8, 9]. Previous experimental conformational analysis investigation on its adsorption on lipid bilayer associated with molecular dynamic simulations in TFE aqueous solution have shown that this distribution of acidic and basic residues and the presence of aspartic acid (D2) at the N-terminus contribute for the helical stabilization of a helical amphipathic structure [10]. There are situations in which bactericide activities of AMPs have to be performed at low or high pH. This distribution of acidic and basic residues can potentially lead this peptide to be pH responsible and desirable to act in these conditions. The pK_*a*_ values of aspartic acids (∼ 4.0) and lysines (∼ 10.4) of MP1 suggest that this peptide most likely would respond to pH changes near these values as observed for another peptide similar to MP1 [11–13]. By substituting the MP1 lysines by histidines, with a lower pK value (*pK_a_* ∼ 6.5), could lead the modified peptide (H-MP1) respond to a narrower pH range roughly two units bellow the physiological one.

In the present manuscript, we have compared the effects of acidic (pH=5.5) and physiological pH solutions on the adsorption of MP1 and H-MP1 to the anionic mixed membrane (7POPC:3POPG) mimicking Gram-negative bacteria plasma membranes [14] and the conformational analyses of these peptides. In this investigation, we have complemented the experimental information of peptide adsorption, insertion into the bilayer, and leakage of lipid vesicles with molecular dynamic (MD) simulations of the peptides in lipid bilayer over to 10 *μ*s each. With this strategy, we got new insights on the atomistic of the peptides’ adsorption process to the anionic membrane. The results indicate that the strong interaction between the AMPs and the bilayer occurs through the N-terminus. Their amphipathic helical structure playing an important role in their positioning into the bilayer.

## 2 Materials and Methods

### 2.1 Chemicals and reagents

1-palmitoyl-oleoyl-sn-glycero-3-phosphocholine (POPC), 1-palmitoyl-2-oeloyl-sn-glycero-3-phosphoglycerol (POPG), 1,2-dioleoylsn-glycero-3-phosphoethanolamine-N-(lissaminerhodamine B sulfonyl) (Rhod-PE) were purchased from Avanti Polar Lipids (Alabaster, AL-USA) and used without further purification. Sodium citrate trisodium salt, sodium borate, mono basic sodium phosphate, sodium chloride (NaCl), calcein were from Sigma Aldrich. Sodium hydroxide (NaOH), sodium fluoride (NaF), hydrochloride acid (HCl), chloroform and methanol were from Merck (Darmstadt, Germany). All these reagents were analytical grade. Ultrapure water Millipore Milli-Q system, (18 MΩ cm) was used for the preparation requiring water. The buffer (CBP) used in the experiments was composed of 1 mM of sodium citrate, sodium borate, and sodium phosphate, at pH 5.5 or 7.4.

### 2.2 Peptide Synthesis

The peptides MP1 (IDWKKLLDAAKQIL-NH2) and H-MP1 (IDWHHLLDAAHQIL-NH2) were prepared by step-wise manual solid-phase synthesis, using N-9-fluorophenylmethoxy-carbonyl (Fmoc) chemistry with Novasyn TGS resin (NOVABIOCHEM, Germany). A side-chain protective group was used t-butoxycarbonyl for lysine. Cleavage of the peptide-resin complexes were performed by treatment with trifluoroacetic acid/1,2 ethanedithiol/anisole/phenol/water (82.5:2.5:5:5:5 by volume), using 10 mL/g of complex at room temperature for 2h. After filtering to remove the resin, anhydrous diethyl ether (SIGMA, USA) at 4 °C was added to the soluble material, causing precipitation of the crude peptide, which was collected as a pellet by centrifugation at 1000 g for 15 min at room temperature. The crude peptides were suspended in water and chromatographed under RP-HPLC using a semi-preparative column (SHISEIDO, Japan; C18, 250 10 mm, 5 *μ*m) at a flow rate of 2 mL/min in the following isocratic conditions with acetonitrile in water (containing 0.1% (v/v) trifluoroacetic acid): 48% (v/v) for MP1, and 46% (v/v) for H-MP1. Under these conditions MP1 was 98% pure, while the purity of H-MP1 was 96 %.

### 2.3 Preparation of large unilamellar vesicles (LUVs)

Lipid films of POPC/POPG mixture at 7:3 molar ratio were obtained from lipids dissolved in chloroform:methanol (2:1) in round-bottom flasks. Organic solvent was removed by drying the lipid film under a N2 flux, and then, under vacuum overnight. Lipid films were then hydrated with CBP buffer containing either 25 mM calcein for leakage experiments or 150 mM NaF for CD or 150 mM NaCl for tryptophan quenching by acrylamide and submitted to an intense vortex. LUVs were obtained by extrusion using an Avanti Mini-Extruder (Alabaster, AL) and double–stacked polycarbonate membrane (Nuclepore Track-etch Membrane, Whatman) in two steps: firstly 6 and 11 times through polycarbonate membranes of 0.4 and 0.1 *μ*m pore size, respectively. Dynamic light scattering measurements with a Zetasizer Nano NS-90 (Malvern Instruments, Worcestershire, U.K.) of pure lipid vesicle suspension showed an average vesicle radius of 66± 6 nm. Only fresh vesicle suspension were used in the experiments. For leakage experiments, gel filtration was used to remove the fluorescent dye outside the liposomes.

### 2.4 Circular Dichroism (CD) Titrations

CD titrations were carried out by adding aliquots of a LUV suspension to a peptide solution at 10.0 *μ*M up to a total lipid concentration of 1.7 mM. After each vesicle addition, the CD spectra were collected over the range 260-190 nm using a spectropolarimeter Jasco J-815 (JASCO International Co. Ltd., Tokyo, Japan) with a Peltier system to control the temperature at 25 °C. These titrations were performed using a 0.2 cm path-length quartz cuvette. The CD spectra were averaged over 10 to 30 scans each one, at a scan speed of 50 nm/min with bandwidth of 1.0 nm, 0.5 s response time and 0.1 nm resolution. Following baseline correction, the observed ellipticity, *θ* (mdeg) was converted to mean residue ellipticity Θ (deg *cm*^2^*/dmol*), using the relationship Θ = 100*θ/*(*lcN*), where */i* is the path length in centimeters, c is millimolar peptide concentration, and N is the number of peptide residues. The *α*-helix fraction (*f_H_*) was evaluated from the observed mean residue ellipticity in 222 nm (Θ_*obs*_) [15]:

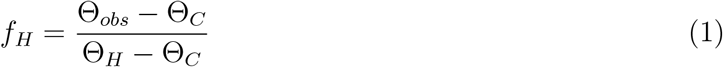

where Θ_*C*_ is molar ellipticity of random coil (Θ_*C*_ = 1500*deg.cm*^2^*/dmol*) and Θ_*H*_ is the molar ellipticity of completely helical peptide, that is given by [15]:

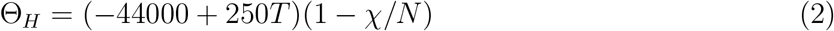

where *χ* = 3 is the number of non-H-bonded CO groups in an amidated peptide, N=14 is peptide length and T = 25 °C. The plots of the observed molar ellipticity in membrane normalized by molar ellipticity of pure peptide (Θ_*obs*_Θ_0_) in 222 nm as a function of the lipid concentration [L] were fitted with the equation [16]:

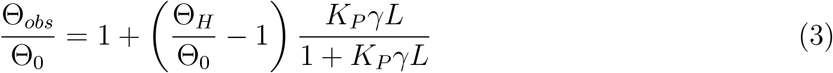

where *K_P_* is the molar fraction partition coefficient and *γ* = 0,75 dm^3^/mol is the lipid molar volume [16, 17].

### 2.5 Tryptophan quenching by acrylamide

Acrylamide fluorescence quenching experiments were performed by adding aliquots of a 2.77 M acrylamide aqueous solution in a, constantly stirred, 5.0 *μ*M peptide solution in 150 mM NaCl solution at desired pH in the absence and in the presence of 500 *μ*M lipid vesicles at 25°C. Tryptophan emission fluorescence spectra were collected from 310 to 450 nm, excited at 285 nm using a PSI spectroluorometer ISS-PC1 (Urbana Champaign, IL, USA). The fluorescence quenching data were analyzed according to the Stern-Volmer equation for collisional quench

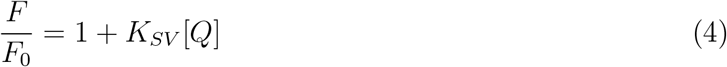

where *F*_0_ and F are fluorescence intensities, measured manually with the cursor at the maximum intensity peak, in the absence and in the presence of quencher respectively. *K_SV_* is Stern-Volmer constant for collisional process and [Q] is the quencher concentration (Lakowickz 1983). In the case of static quenching the equation used for the numerical fitting was:

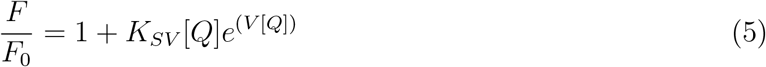

where V is a static quenching constant[18].

### 2.6 Lytic Activity in LUVs

Dye leakage from the LUVs: Calcein leakage from LUVs was monitored by fluorescence spectroscopy by measuring the fluorescence emission intensities at 520 nm (excited at 490 nm) using an ISS PC1 spectrofluorometer (Urbana Champaign, IL, USA). The percentage of calcein leakage induced by the peptide was calculated with equation:

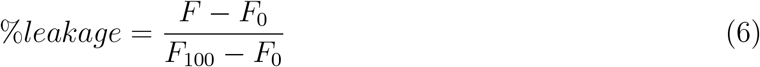

where F is the fluorescence intensity obtained after the vesicle/peptide interaction, F_0_ is the fluorescence intensities in the absence of peptide and F_100_ correspond to 100% of leakage, as determined by the addition of 20 *μ*l of 10% Triton X-100 solution.

### 2.7 Molecular Dynamics Simulations

To perform the Molecular Dynamics simulations of the investigated systems we build a lipid bilayer composed by 120 phospholipids (60 on each leaflet) of POPC and POPG in a 70:30 ratio with the Membrane Builder plugin [19] from CHARMM-GUI website [20] using CHARMM 36 force field [21]. The bilayer was solvated on a TIP3P [22] water box of about 65×65×120 Å and the water molecules located at the hydrophobic region were removed. Enough Na^+^ ions were added to neutralize the system, and 150 mM of NaCl were added to mimic the physiological environment. We used VDM [23] to perform all these procedures. This simulation box was equilibrated with 10000 steps of conjugate gradient energy minimization and 100 ns of equilibrium MD using the software NAMD [24].

The peptides were generated in *α*-helix configuration from their amino acid sequence using Molefacture plugin of VMD and CHARMM 36 force field. For both peptides the C-terminus is amidated and the N-terminus is positively charged. Each one was inserted on the water phase of the previously equilibrated hydrated bilayer simulation box about 10 Å far from the bilayer surface. It is noteworthy that we correct the number of ions after including the peptide on this box, represented on figure 1. The charges for both peptides were chosen based on the pK_*a*_ of each titratable residue and the pH of the simulation. The new systems with the bilayer and one peptide in aqueous solutions were equilibrated with 10000 steps of energy minimization and 10 ns of molecular dynamics, with position restraints on the peptide backbone for solvent relaxation, using NAMD.

**Figure 1:**
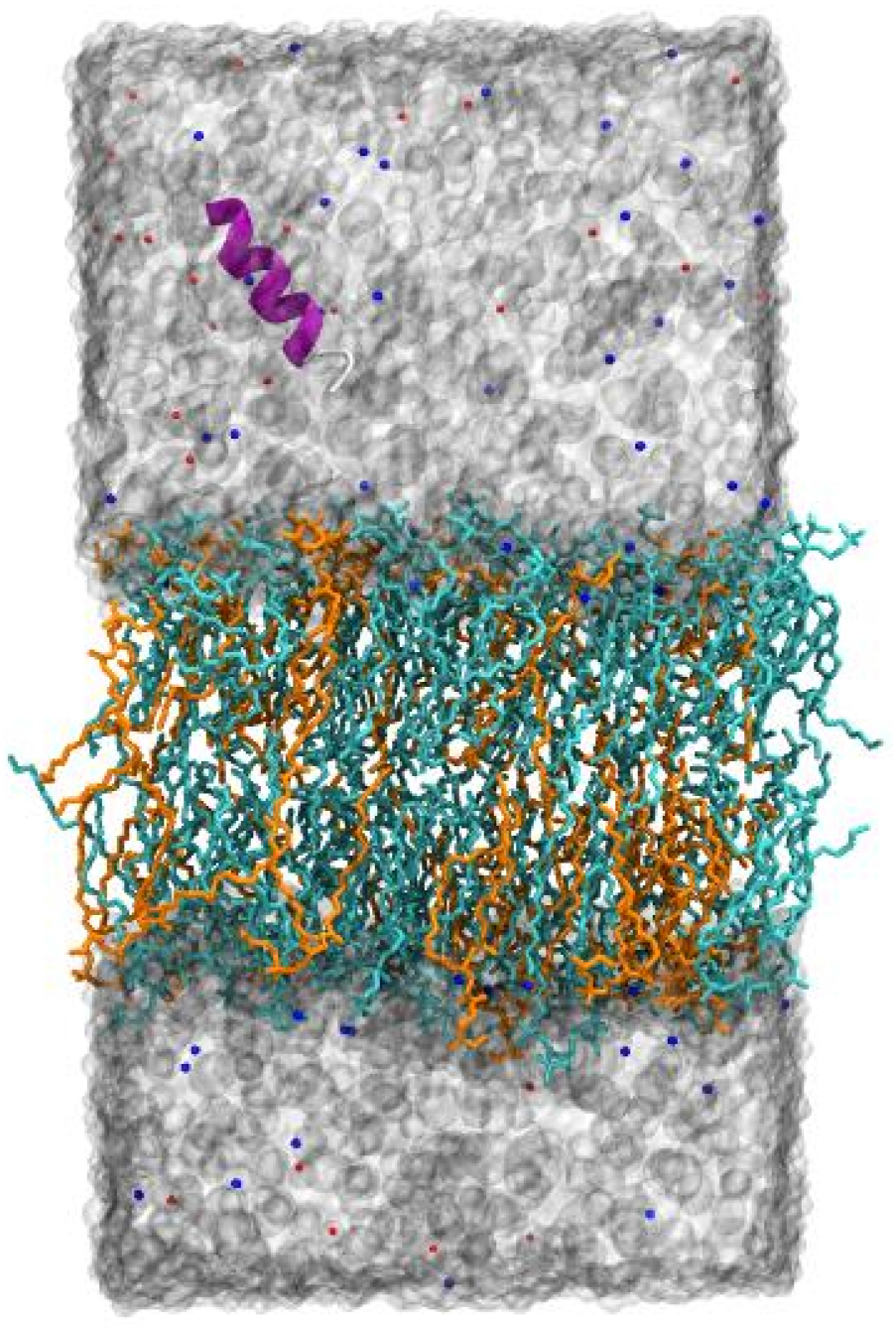
Representation of the simulation box containing water (gray surface), the bilayer of POPC (cyan) and POPG (orange), ions *Na*^+^ (blue) and *Cl^−^* (red) and the peptide on secondary structure representation (*α*-helix is represented on purple and coil on white). At the beginning of the simulation the peptide is randomly placed at about 10 Å of the lipid bilayer.

All the MD simulations were performed with a 2 fs time-step, periodic boundary conditions, on the NPT ensemble at 298 K and 1 atm. Temperature and pressure were modulated by Langevin thermostat[25] and Langevin piston[26]. The SHAKE algorithm [27] was used to control the lengths of atomic bonds, and to preserve the geometry of water molecules the SETTLE algorithm [28] was applied. The Van der Waals interactions were calculated with a cut off of 12 Å with switching distance of 14 Å and the long range interaction were treated using Particle Mesh Ewald (PME) method[29].

At this study, three systems were investigated, one for MP1 because the pH range (5.5 to 7.4) is far from the *pK_a_* of the titratable residues, and two situations for H-MP1, one at pH 7.4 and other for pH 5.5. The MP1 protonation state used in these MD simulations were N-terminus and lysines protonated and aspartic acids deprotonated, for the H-MP1 at pH 7.4 the N-terminus was protonated and histidines and aspartic acids were deprotonated and for H-MP1 at pH 5.5 the N-terminus and histidines were protonated and aspartic acids were deprotonated.

To better sample the system’s phase space, twelve simulations with different initial velocities were performed for each system (MP1, H-MP1 protonated and, H-MP1 deprotonated), since a recent work indicates that the use of several replicas is often more accurate than only one long simulation[30]. Overall, around 2 *μ*s of Molecular Dynamics were performed for H-MP1 deprotonated, and 10 *μ*s each for MP1 and H-MP1 protonated. To detect adsorption events, we measured the distances between the N-terminus, C-terminus, and the center of mass of the peptide and the center of mass of the bilayer for all the simulations. We observed adsorption only in systems where the peptides are protonated and we extended the simulation time for the ones these distances were lowest. It was not observed an insertion in the bilayer of the H-MP1 deprotonated on any simulation and that is why they were not extended as the other systems. The absence of adsorption events for this system is expected since both, peptide and bilayer, present net charge with the same polarity. This does not agree with experimental results and it is likely due to the fixed charge model used in our MD simulation. Probably, charge regulation is important for this adsorption process and its proper modeling should be done with variable charge techniques as Constant pH Molecular Dynamics (CpHMD).

The analysis were made using the VMD software [23] and some of its plugins like Namdenergy, Membplugin [31] and Stride [32] to calculate the secondary struture.

## 3 Results and Discussion

### 3.1 The pH effect on the secondary structure and on the affinity of MP1 and analog H-MP1 to POPC/ POPG (70/30) LUVs

The experimental conformational analyses of MP1 and H-MP1 at acidic and neutral pH in the absence and the presence of vesicles were performed by CD spectroscopy. Fig. 2a shows CD spectra of H-MP1 and MP1, at acidic pH and fig. SM1 at neutral pH, respectively in the absence and the presence of lipid vesicles at [L]/[P]=30. At acidic and neutral solution pH (5.5 and 7.4) and in the absence of lipid vesicles, H-MP1 and MP1 CD spectra show an intense negative band near 200 nm suggesting the secondary structure was absent. The MP1 spectra at both solution pH show a negative drift from the baseline at 220 nm with a mean molar ellipticity roughly three times larger than for the analog H-MP1 at pH 5.5. This drift also suggests that the peptides had a small amount (∼ 10 to 30%) of a helical structure at acidic pH for analog and MP1 respectively. The addition of lipids at [L]/[P]=30 spectra shape changed with two negative dichroic bands at 220 and 208 nm and a positive one near 190 nm, characterizing an *α*-helix structure. The intensities of these negative bands increased with the [L]/[P] ratio. At the same total lipid concentration, the CD spectra of H-MP1 showed more intense bands (222 and 208 nm) at pH 5.5 compared to the neutral solution, while MP1 they were less sensitive to the pH. For both peptides, the intensity of the 208 nm band exceeded that 222 nm indicating that the adsorbed peptides were in monomeric form [33, 34]. Titrating the peptide solutions with 7POPC:3POPG vesicles suspension, up to the spectrum saturation, allowed assessing the effects of the solution pH on the affinities of both peptides to this lipid bilayer. Throughout the titration, an iso-dichroic point around 204 nm was indicative of equilibrium between two peptide states: free and adsorbed. The intensities of the negative bands increased with the total lipid concentration up to a limit. The insets of Fig. 2a and SM1-a shows the adsorption isotherms, the dependence of the normalized molar ellipticity on the total lipid concentration for H-MP1 and MP1 respectively at pH 5.5 and 7.5. These isotherms were analyzed using a Langmuir type isotherm whose best fitting are continuous lines in this figure. Adsorption constants (Kp) and the maximum helical content (fH), obtained from the numerical fit of these isotherms, are displayed in the table I. These results show that the solution pH strongly modulated the affinity of the analog H-MP1 to 7PC:3PG vesicles and MP1 was almost insensitive to the solution pH. The adsorption of the analog H-MP1 decreased with the increase of solution pH. At acid solution, the analog H-MP1 partitioned to these vesicles with a constant that was about 20 % larger than that of MP1. At neutral pH, the analog partition constant was roughly three times smaller compared with MP1.

**Figure 2:**
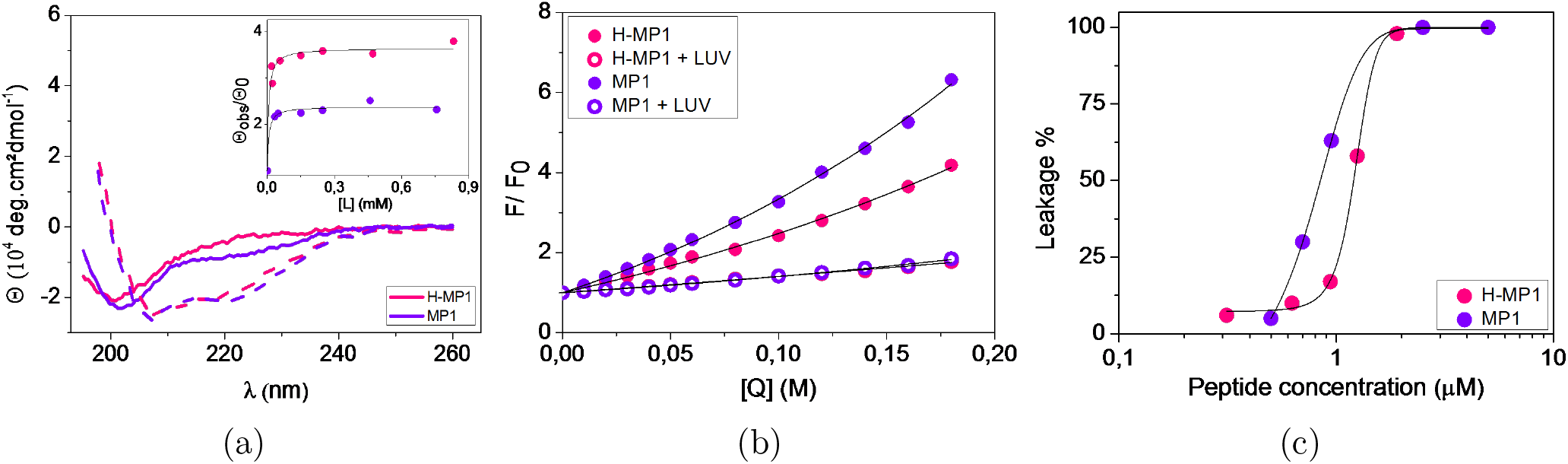
(a)-CD spectra of 10*μ*M of H-MP1 (magenta) and MP1 (purple) at pH 5.5, 25 °C in buffer (line) and in (7POPC:3POPG) LUVs [L]/[P]=30 (trace). Inset CD adsorption isotherms:*θ*/*θ*_0_ vs total lipid concentration [L] at pH 5.5. Lines are best fitting with Lagmuir model eq. 3 (b)-Stern-Volmer plots for the tryptophan quenching by acrylamide of H-MP1 (magenta) and MP1 (purple) at pH 5.5, in buffer (closed) and in LUVs (open circles). lines are best fitting using eq.5 and 4 respectively (c) Dose-response curves, % Calcein leakage from LUVs (100*μ* M lipid) induced by H-MP1 (magenta) and MP1 (purple) at pH 5.5 and 25 °C after 10 min. contact.

**Table 1:** Summary of experimental results: Partition constants K_*p*_, *α*-helix fraction f_*H*_ and ratio of Stern-Volmer constants K_*SV V*_ /K_*SV B*_ at pH 5.5 and 7.4

| . SD from three different experiments. |  |  |  |  |  |
| --- | --- | --- | --- | --- | --- |
| Peptide | pH | $K_p \times 10^3 (M^{-1})$ | $f_H$ | $K_{SVV}/K_{SVB}$ | EC<sub>50</sub> ($\mu$ M) |
| MP1 | 5.5 | $47 \pm 18$ | $0.74 \pm 0.02$ | $0.19 \pm 0.03$ | $0.8 \pm 0.1$ |
| | 7.4 | $87 \pm 13$ | $0.65 \pm 0.04$ | $0.20 \pm 0.01$ | $0.7 \pm 0.1$ |
| H-MP1 | 5.5 | $119 \pm 35$ | $0.68 \pm 0.02$ | $0.32 \pm 0.02$ | $1.2 \pm 0.1$ |
| | 7.4 | $37 \pm 13$ | $0.46 \pm 0.03$ | $0.36 \pm 0.01$ | $7.6 \pm 0.6$ |

### 3.2 The insertion of peptide into the lipid bilayer

Lytic peptide adsorption on lipid bilayer is, in general, accompanied by its insertion into the hydrophobic phase of the bilayer. Tryptophan fluorescence quenching by acrylamide allowed assessing peptide insertion into the bilayer. Fig. 2b shows the dependence of the normalized tryptophan fluorescence intensities as a function of acrylamide concentration at pH 5.5 and Fig. SM2 the same plots at neutral pH. These plots were analyzed using the Stern-Volmer model for collisional quenching and are the continuous lines in Fig. 2b and Fig. SM2 are the best fit obtained using eq.5 and 4 respectively in buffer and in vesicles providing the Stern-Volmer constants K_*SV*_. The ratios of the Stern-Volmer constants, in vesicle K_*SV V*_ and buffer K_*SV B*_ allowed estimating how the tryptophan is screened from the quencher and consequently from solvent. The smaller this ratio more protected from the solvent is the tryptophan and most likely more deeply inserted into the hydrophobic phase of the membrane. Table-I displays the values of the K_*SV V*_ /K_*SV B*_ ratios indicating that for MP1 this ratio was insensitive to the solution pH, while for H-MP1 it was roughly 30 and 50% larger compared to MP1. MP1, most likely, inserted more deeply into POPC/POPG bilayer in comparison to the analog.

### 3.3 pH Effect on the Lytic Activity in LUVs

The lytic activity of both peptides in LUVs was investigated by leakage experiments using calcein entrapped LUVs. When inside the liposome, the calcein fluorescence is self-quenched. Upon peptide action, calcein is released, recovering the fluorescence intensity. The time course of fluorescence recovery for both peptides in different peptide concentrations allowed obtaining dose-response curves showed in Fig.2c and Fig. SM3 at acidic and neutral pH, after 10 min. of peptide action. These dose-response curves indicated that both peptides achieved complete leakage irrespective of the solution pH. The leakage efficiency, measured by EC50, the peptide concentration to induce 50% of calcein release, showed that MP1 was more efficient compared to the analog and that MP1 efficiency (EC50 0.8 *μ*M) was roughly independent of the solution pH. Solution pH strongly modulated the lytic efficiency of the analog. At acidic pH, H-MP1 was almost seven-times more efficient (EC50 1.2 *μ*M) than in neutral pH (EC50 8*μ*M). In this way, the lytic activity in liposomes indicated that the substitution of lysines per histidines in MP1 resulted in a high pH-sensitive peptide.

### 3.4 Adsorption of MP1 and H-MP1 by Molecular Dynamics Simulations

The precedent experimental analysis of the effects of solution pH on the conformation, adsorption and insertion of the peptides in POPC:POPG (7:3) lipid vesicles should be understood only as an average behavior of system peptide/lipid vesicle devoid of molecular and atomistic details. Aiming to gain new insights on these levels the experimental data were complemented with Molecular Dynamics simulations. In these simulations the peptides were initially in a helical conformation and at 10 Å from the phosphorous of the lipid bilayer as shown in Fig. 1.

#### 3.4.1 Secondary structure analysis

As we already know from CD experiments, that the peptides’ adsorption to the lipid bilayer is accompanied by an increase of *α*-helix secondary structure of both peptides, this conformational change was also observed on the MD simulations. In order to save computational time in these simulations the peptides already started as an *α*-helix that is previously known to be the predominant structure (figure 2a). In the absence of lipid bilayer 1 *μ*s MD simulations was performed for each peptide in 150 mM of NaCl aqueous solution. The results of these simulations are presented on Supplementary Material. Fig. SM4a and SM4b shows the residues in helical conformation, calculated with STRIDE [32], for the two peptides as a function of the simulation time. These figures show that the peptides significantly loose their helical conformation in aqueous solution. H-MP1 quickly loose half of helical conformation up to 200 ns and then completely loose all the initial conformation. MP1, on the other hand, loose 20% of helical conformation in the first 200 ns and after 600 ns remained partially at the helical conformation roughly 50% within its central residues. These results are in reasonable accordance with those obtained from the CD spectra in pure buffer with roughly 10 and 30 % of helical structure for H-MP1 and MP1 respectively.

STRIDE consider to classify the type of secondary structure, besides the backbone hydrogen bonds, the *ϕ* and *ψ* dihedral angles. Using the software DSSP [35], that not considers the dihedral angles, we obtained one residue less on *α*-helix configuration. Both the softwares are good to classify the secondary structure on proteins, but they can overestimate the *α*-helix content on small peptides [36]. To gain a better understanding about the time evolution of helical content, persistence of the hydrogen bonds in the backbone of the peptide between residues *i* and *i*+4 were also analyzed. Fig. SM5 shows the lifetime of backbone hydrogen bonds between the carbonyl of each MP1 residue *i* and the amino group of the residue *i* + 4. This figure shows that only five residues are involved in H-bonds during at least 50% of the simulation time, indicating roughly 36% of stable helical structure, in agreement with the experimental results. The life-times of the backbone H-bonds for H-MP1, also shown in the Fig. SM5, indicates that H-bonds were persistent for only 20% of simulation time and, consequently, the helical content is very low which is also in good accordance to the experimental results.

An important structural analysis to investigate the role of side chains on the *α*-helix stabilization is to look for the distance of the ion-pairs or salt bridges formed by the charged residues. Salt bridges are long range and non-directional interactions, so they can be formed when the structure is on random coil, while backbone’s hydrogen bonds are small range and directional interactions. In alanine based peptides, salt bridges help to maintain the charged residues closer, favoring the formation of *i*-*i* + 4 hydrogen bonds on the backbone that are necessary for the peptide to acquire an *α*-helix conformation [37–39]. For MP1 in 150 mM NaCl aqueous solution, represented on Figure SM7a, one can see that the distances between the lysine (LYS4) and the aspartic acids, ASP2 and ASP8, are compatible with the formation of two salt bridges that seems to commute between them during the simulation time with a higher prevalence for ASP8-LYS4. On the other hand, the distance between the aspartic acids and histidines in H-MP1, shown in the Fig. SM7b, were larger than those observed for MP1 suggesting that salt bridges were hardly formed, what can explain the low helix conformation of this peptide in aqueous solution.

As expected from the CD spectra, upon the peptides’ adsorption to the bilayer the amount of helical structure increases significantly. Figures SM6a and SM6b display the helical conformation per residue along the 1.2 *μ*s simulation time for MP1 and H-MP1 in the presence of the bilayer described on section 2.6. These figures show that for MP1, after the adsorption (200 ns), 13 residues were in *α*-helix conformation leading to 93% of helical content, well above the experimental result obtained from CD. The H-MP1 peptide loses most of its initial *α*-helix secondary structure in the first 400 ns. After this simulation time, when the H-MP1 completely adsorbs (see next section), the peptide recovers the lost helical content, displaying 11 residues in this secondary structure corresponding to 79% of its total length, a percentage, again, higher than the experimental findings.

The calculation of lifetime percentage of backbone *i*-*i* + 4 H-bonds improved our secondary structure estimation (Figure 3). The backbone hydrogen bonds for adsorbed MP1 peptide between the pairs of residue 1-4 to 7-11 were prevalent for at least 50% of the simulation time, corresponding to 7 of 10 possibilities, or 8 residues, that most likely should be engaged in helical structure. Comparing these results to those shown in the Fig. SM5 of the supplementary material, that presents the same type of hydrogen bonds for the simulation on bulk, the MP1 has 5 hydrogen bonds (2-5 to 6-10) that exists for more than 50% of the simulation time indicating an increase of helical conformation upon peptide adsorption to the bilayer. These results also show the C-terminus region is, mostly of time, on a non-helical conformation, assuming a random coil secondary structure as shown in Figure SM6.

**Figure 3:**
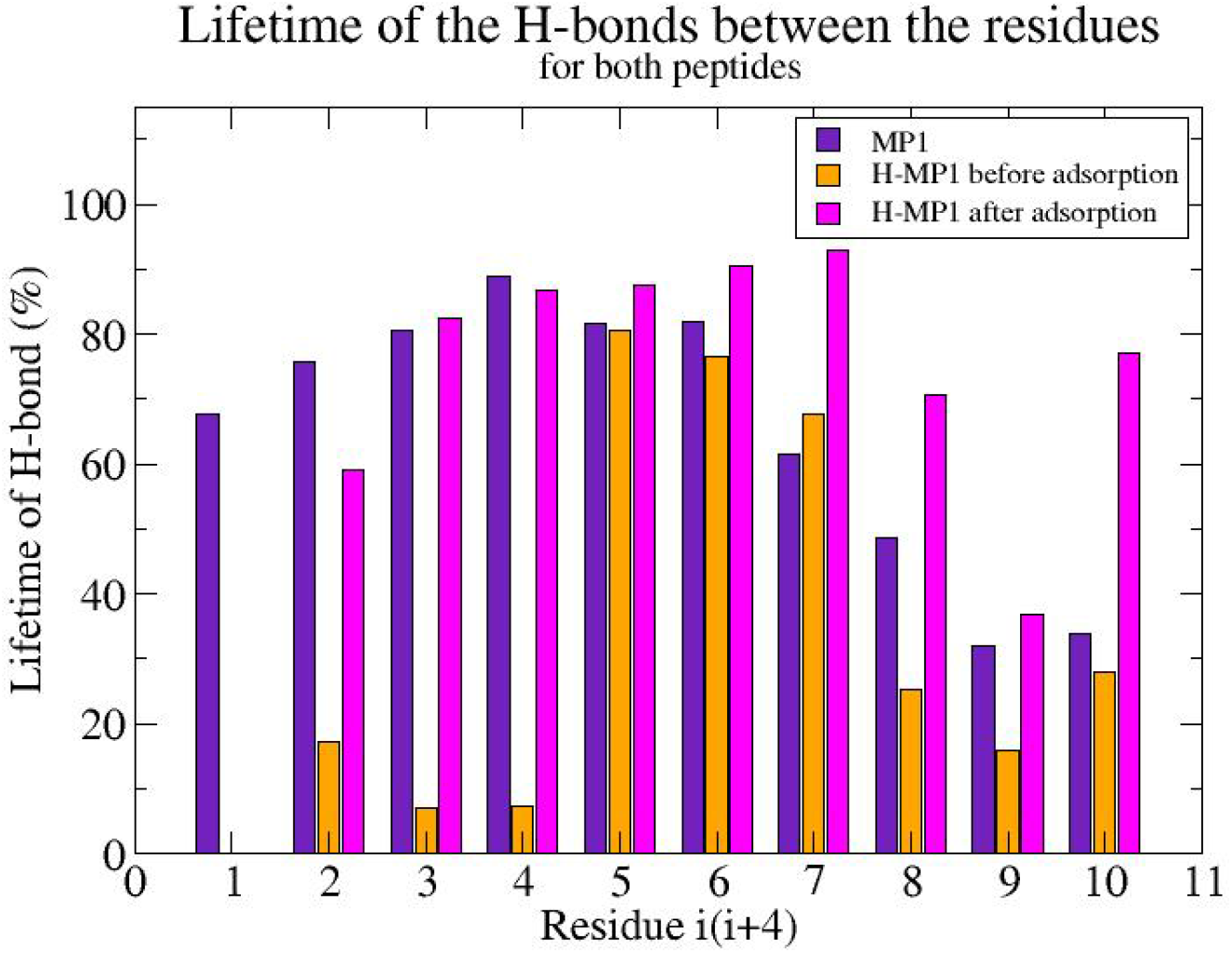
Percentage lifetime of backbone hydrogen bonds between residues *i*-*i* + 4 for MP1 and H-MP1 to investigate the *α*-helix stability during the simulation. The MP1 is represented on purple and the H-MP1 is divided on two situations: before the adsorption on orange, and after on magenta.

The *i*-*i* + 4 H-bonds profile for H-MP1 shows different behavior before and after the complete peptide adsorption. At the simulation’s beginning, when only the N-terminus is adsorbed and the peptide is perpendicular to the bilayers plane (see Figure 5b), the helical content decreases and the H-bonds stable for more than 50% of the time fall from ten to three (orange bars in Figure 3). This is an intermediate conformation observed between the peptide in 150 mM NaCl aqueous solution (Figure SM4), and its conformation after 400 ns, when the whole peptide adsorbs and 8 out of 10 *i*-*i* + 4 H-bonds are stable most of the time (magenta bars in Figure 3).

As far as we know, this is the first time the complete folding process for an MP1 analog is observed at atomic resolution. From our simulations, it is possible to observe in details the peptide’s secondary structure change from the coil rich unfolded state at bulk, the intermediate state when the N-terminus adsorbs and the peptide starts to feel the influence of the bilayer over its secondary structures, and the folded state when the peptide is fully adsorbed, parallel to the bilayer’s plane, in an amphipatic helical rich conformation induced by the hydrophilic/hydrophobic interface provided by the bilayer.

Concerning salt-bridges, it is noteworthy that in the adsorption simulation of MP1, the LYS4 display a stable salt bridge with ASP8 (Figure SM8a), as also observed in the aqueous solution simulations of this peptide. For H-MP1, after the N-terminus adsorption at the simulation’s beginning, the distance between ASP2 and HSP5 (Figure SM8b) indicates a salt bridge that probably sustain the *α*-helix portion at the middle region. After 400 ns, when the whole peptide adsorbs, another salt bridge is formed between ASP8 and HSP4 what possible helps the peptide’s folding.

#### 3.4.2 Adsorption Process

MD simulations provide useful information at atomic resolution of the adsorption processes of both peptides to this anionic mixed (7POPC:3POPG) bilayer. The adsorption of both peptides starts by inserting their charged N-terminus portion into the bilayer as noticed from the figures 4. After this first contact, the system evolves as the peptide’s middle region and amidated C-terminus also adsorbs and the final peptide orientation is parallel to the bilayer plane, tilted slightly due to the burial of C-terminus into the hydrophobic region (Figures 4 and 5). It is noteworthy that, when adsorbed, the tryptophan residue of both peptides lies on the hydrophilic-hydrophobic interface of the bilayer, what was expected considering the fluorescence quenching experiment (Figure 2b).

**Figure 4:**
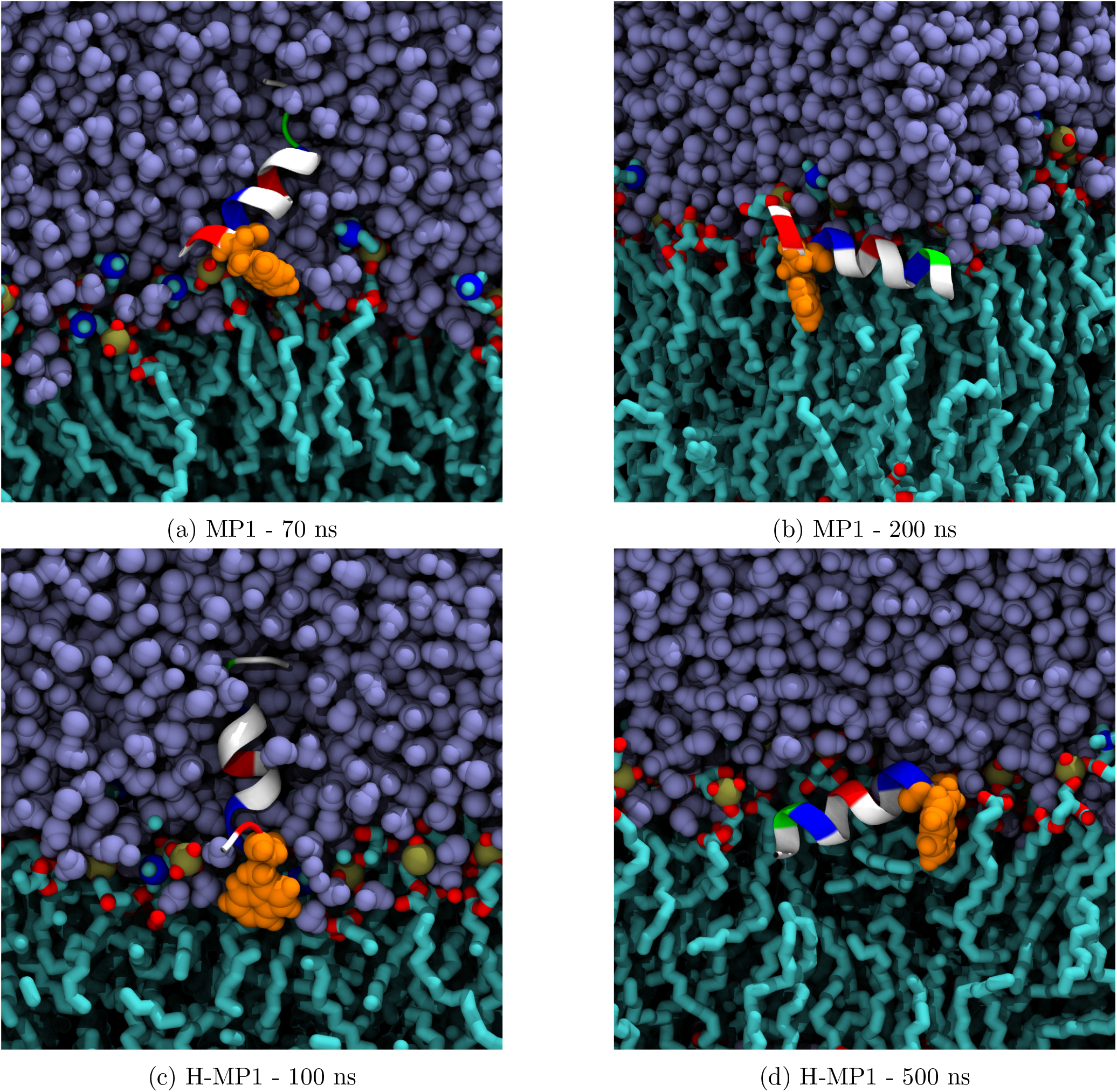
Snapshot showing how the MP1 and H-MP1 adsorption occurred. The MP1 adsorption’s figures right before and after the adsorption are represented on figures 4a and 4b, respectively, and the H-MP1 before and after the adsorption is represented on figures 4c and 4d, respectively. The peptides are represented by its secondary structure and colored by the residue type (hydrophobic residues represented on white, positive ones on blue, negative ones on red and polar uncharged on green), the TRP residue represented on orange, water on ice blue, and lipids by its atom name, cyan being carbon, nitrogen on blue, oxygen on red and phosphorus on gold.

**Figure 5:**
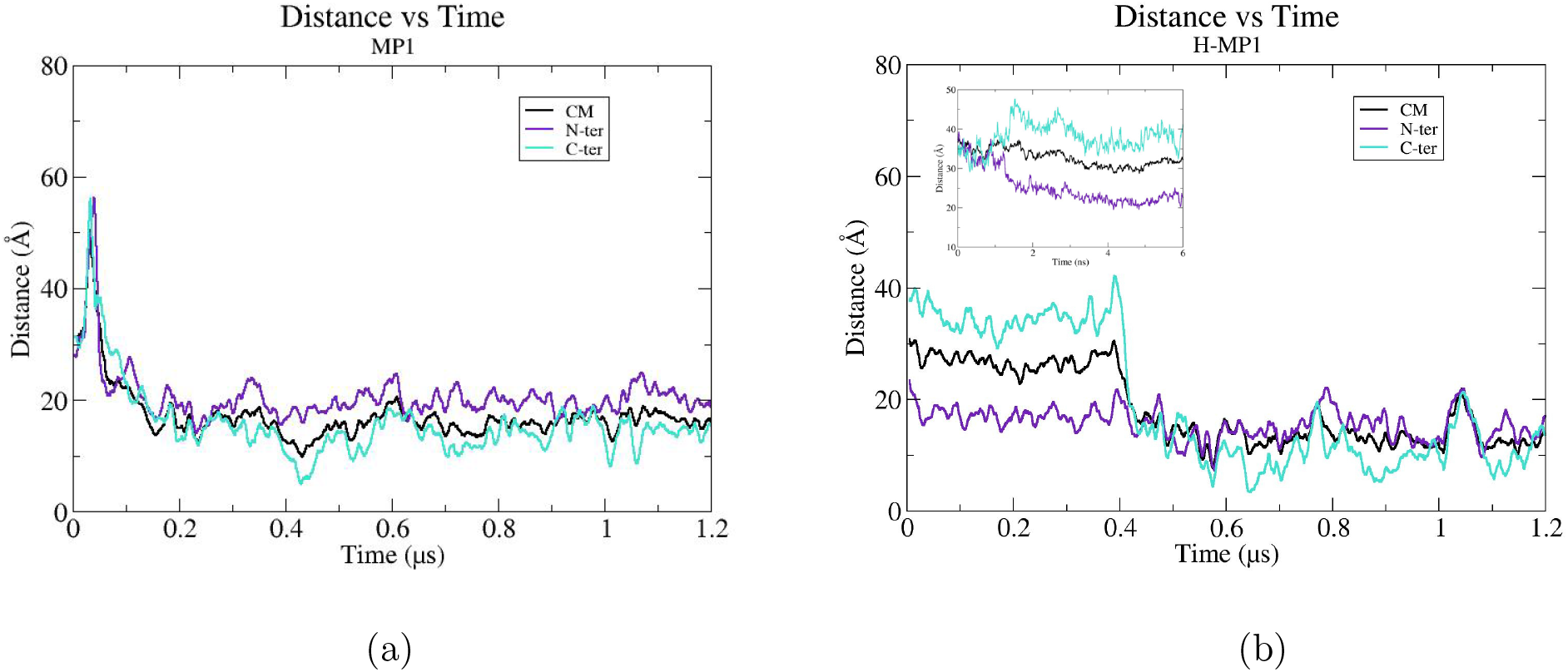
Distance between different regions of the peptides and the center of mass of the lipid bilayer. Three positions for both peptides were considered here, the center of mass (CM) considering the entire peptide on black, the N-terminus being the center of mass of the first residue (ILE1) on purple, the C-terminus being the center of mass of the last residue (LEU14) on cyan. (a): MP1, (b): H-MP1, inset: The first 6 nanoseconds of the simulation when the peptide is outside the bilayer (more than 20 Å of distance), we can see that the peptide started at a parallel position to the bilayer and quickly moved to a perpendicular position and started to approach.

The orientation of each peptide relative to the lipid bilayer throughout the simulation time was assessed by measuring the distances of its N- and C-terminus and of its center of mass (CM) to the bilayer CM at each simulation time. These distances, displayed in the figure 5a for MP1, shows that the adsorption occurred by approaching its N-terminus to the bilayer within 70 ns, then at 180 ns almost the entire peptide is inserted into the bilayer parallel to its surface (XY plane). For H-MP1, shown in fig. 5b, at 6 ns of the simulation the N-terminus approaches the bilayer and begins to invade the hydrophilic region until 50 ns of simulation. The complete adsorption occurred after 400 ns with the same orientation to the bilayer as MP1. The distances of CM (black line), C-terminus (cyan line) and N-terminus (purple lines) to the bilayer CM before 400 ns show the peptide is oriented perpendicular to de bilayer. After 400 ns of simulation these distances were roughly the same during the rest of simulation, indicating that the peptide re-oriented itself to be parallel to the bilayer similar to the orientation of MP1. These results showed that the interaction between the the N-terminus positive charge and the phospholipids’ phosphate groups seems to be the leading factor driving the adsorption process for both peptides. However, as we can see on figure 5b, after the adsorption, the C-terminus of the peptide is more inserted on the bilayer, probably because of the amidation of the C-terminus and the predominance of hydrophobic residues. It was also observed that once adsorbed, the peptides stay in this orientation until the end of the simulation.

Although the distance between the peptide and the center of mass of the bilayer provided us information about the adsorption’s dynamic, it was also worthy to investigate the environment where the peptide is inserted into the bilayer. This was assessed by the average number density profile for the bilayer, water and, antimicrobial peptide before and after its adsorption (Figures SM9). These results indicate that the peptide density profile moved from the outside to the inside of the bilayer, being completely inserted in the hydrophilic/hydrophobic interface region where water and phospholipids coexist. The amino acids orientations of the adsorbed peptides at the hydrophilic/hydrophobic interface are indicated in figure 4. These figures show that the peptides’ hydrophobic faces are buried in the acyl tails region of the bilayer while the hydrophilic residues are interacting with water and both the phospholipids’ polar heads.

The peptides’ positioning on the bilayer (Figure 4) were experimentally corroborated correlating the average position of its tryptophan residue with the ratio of the Stern-Volmer constants (K_*SV V*_ /K_*SV B*_) obtained by the tryptophan fluorescence quenching by acrylamide. The frequency histograms of figure 6 were made measuring the distances between center of masses of TRP3 and the bilayer CM throughout the simulation time. The most frequent distance obtained for MP1 is a narrow peak centered between 11.3 and 14.7 Å, and for H-MP1 a wider peak between 12.9 and 16.5 Å. From these results we can conclude that the TRP residue lies on the hydrophilic/hydrophobic interface of the bilayer, as shown on figure 4 and also indicated by the experiment. This arrangement was already expected since the indole bicyclic structure of tryptophan side chain presents hydrophobic and hydrophilic chemical groups. From the Molecular Dynamics we can also note that the tryptophan residue of MP1 is slightly more buried into the bilayer hydrophobic phase compared to the H-MP1 TRP, in good accordance with fluorescence quenching results (figure 2b).

**Figure 6:**
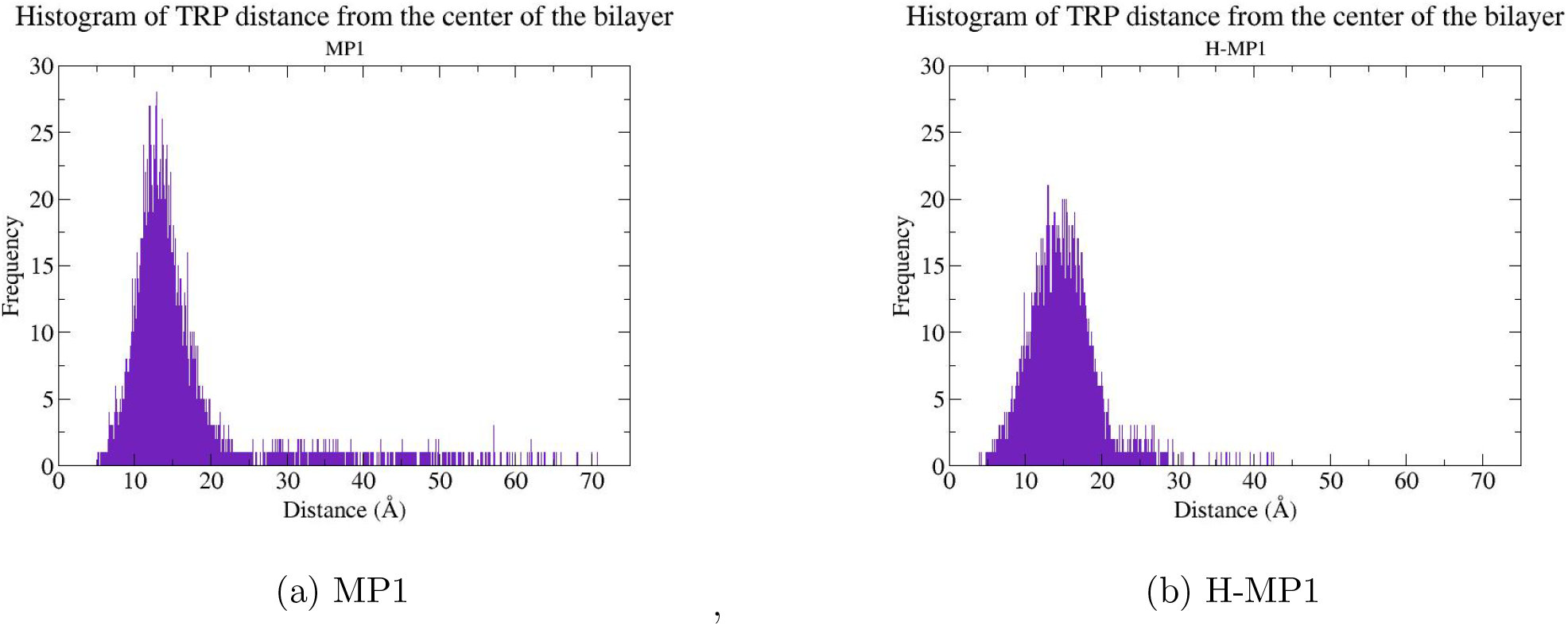
Histogram of the distance between the center of mass of the tryptophan and the center of mass of the bilayer throughout the simulation, showing what is the most frequent distance between this amino acid and the bilayer’s CM.

#### 3.4.3 Energy analysis

Another important information provided by the MD simulations regards the detailed interactions of the peptides’ residues with the phospholipids. Electrostatic and Van der Waals interaction energies were assessed by calculating via Namd energy plugin on VMD the non-bonded interaction energies between each amino acid and the whole lipid bilayer for every frame of the simulation, then, it was calculated the average and standard deviation on the last 500 ns of simulation, when the peptides were fully adsorbed. The obtained results are shown on figures 7a and 7b for each amino acid.

**Figure 7:**
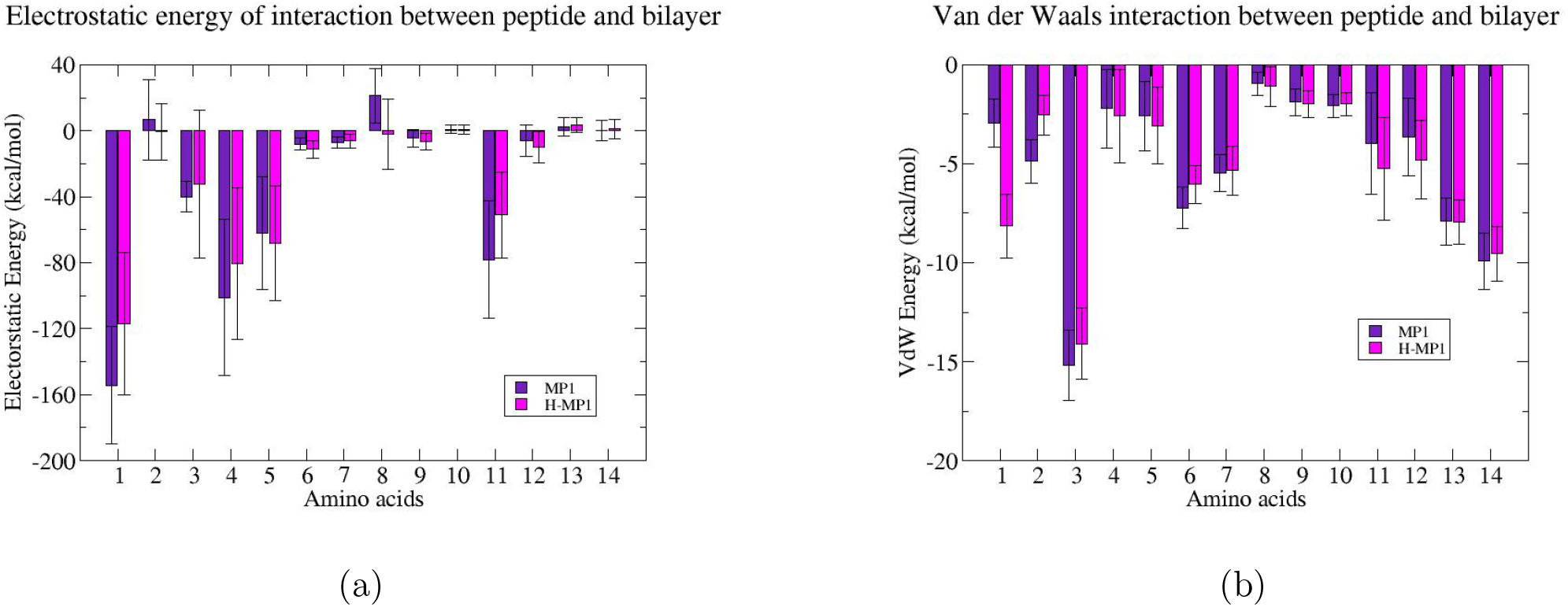
Electrostatic energy and VdW interaction energy between each amino acid of the MP1 and H-MP1 peptides and the lipids of the bilayer and its correspondent standard deviation. MP1 is represented on purple and H-MP1 on magenta.

The Fig. 7a shows that the strongest electrostatic interaction occurs between the positive charge of the N-termini amino-groups and the bilayer, most likely, the phosphate groups. This interaction is slightly more pronounced for MP1 compared to the analog. Lysines 4 and 5 contributes roughly with half the energy of N-terminus and were similar to that of the LYS 11. The two acidic residues (ASP2 and ASP8) showed negligible or small unfavorable electrostatic interactions with the bilayer. The peptide’s orientation places these residues in a position where their attractive interaction with the choline groups are partially offset by their repulsive interaction with the phosphate groups. Excepting for HIS5 all the H-MP1 residues showed smaller electrostatic energies compared with MP1 and for HIS5 they were roughly the same. The most favorable Van der Waals interaction energies were at least 10 times less intense compared with the strongest electrostatic interaction (N-terminus amino group). TRP3 had shown the most pronounced VDW contribution that was similar for both peptides. Although TRP3 have the most favorable VDW energy (∼ −14 kcal/mol), it was less than half compared to its electrostatic energy (∼ −40 kcal/mol) reinforcing the contact of this residue at hydrophobic/hydrophilic bilayer interface. The VDW energies of other non-polar residues LEU6 and LEU7, positioned inside the hydrophobic region of the bilayer, were well bellow that of TRP3 while ILE13 and LEU14 in C-termini were roughly half of the tryptophan for both peptides. The ASP2 unfavorable electrostatic energy has almost the same value of VDW component, suggesting that this energy could also be compensating the unfavorable one. Although the difficulty in estimating the electrostatic and VDW energetic contributions to the adsorption process, the comparison of the partition constants of H-MP1 bearing different net charges (at pH 5.5 and 7.5) evidences the dominance of the electrostatic interactions in the peptides adsorption and in maintaining the stability of the adsorbed peptide into the bilayer.

It is worthy to notice that the ionization state of each amino acid is the same through the simulation, which is a limitation of the traditional Molecular Dynamics method. Constant pH simulations are being made for these systems to provide a better description of the role of charge regulation in the adsorption process and will be the subject of future studies.

## 4 Conclusion

In the present, we investigated, both experimentally and by MD simulations, the solution pH influences on adsorption of two peptides to anionic mixed bilayer and their spontaneous insertion into the hydrophobic phase of the lipid bilayer. The results showed that the analog H-MP1, with histidines substituting the four lysines of MP1, lead to a more selective response to the solution pH in comparison to the parent peptide. Both peptides presented similar behaviors when interacting with the bilayer, increasing the helical content when adsorbed parallel with the bilayer plane. Although both peptides showed similar final adsorbed states, the processes involved in their adsorptions were different. Prior adsorption peptides landed the bilayer touching it with their N-termini. MP1 then quickly lays on the bilayer, the analog remains perpendicular to the bilayer for roughly 400 ns, losing most of its *α*-helix secondary structure, and then suddenly got also parallel to the bilayer, recovering its original helical content due to the influence of the hydrophilic/hydrophobic interface environment. The amount of helical structure determined from MD simulations calculated using *i*-*i* + 4 backbone H-bonds was in better accordance with those determined from CD spectra than those estimated using package STRIDE, which seems to have overestimated the helical amount for the peptides. MD simulations evidenced that both in bulk and bilayer peptides salt bridges between their lateral chain contributed to stabilizing the secondary structure. Overall, both peptides inserted into and most likely disturbed the bilayer lipid packing that is probably important for the lytic activity. The peptides adsorbed states were strongly favored by electrostatic and by van der Waals interactions to a lesser extent.

## 5 Acknowledgements

The authors acknowledges financial support from Brazilian agencies: São Paulo Research Foundation-FAPESP (JRN: FAPESP grant # 2015/25619-9; MSP grant# 2016/16212-5 and ASA FAPESP grant # 2010/18169-3. IBSM PhD grant CAPES, TGV PhD grant CNPq. MSP CNPq/INCT iiii. JRN and MSP are researchers for Brazilian Counsil for Scientific and Technological Development (CNPq). This research was supported by resources supplied by the Center for Scientific Computing (NCC/GridUNESP) of the São Paulo State University (UNESP), the “Centro Nacional de Processamento de Alto Desempenho em São Paulo (CENAPAD-SP)” and also for the National Laboratory for Scientific Computing (LNCC/MCTI, Brazil) for providing HPC resources of the SDumont supercomputer, which have contributed to the research results reported within this paper. URL: http://sdumont.lncc.br.

## Author contributions

IBSM and ASA performed the simulations and computational analysis. TGV and JRN perfomed the experiments and experimental analysis, MSP and BMS synthesized and purified the peptides. All authors discussed the results and participated on writing and reviewing this paper.

## Notes

The authors declare no competing financial interest.

