## Supplementary material for "The effect of acidic pH on the interaction and lytic activity of MP1 and its H-MP1 analog in anionic lipid membrane: a biophysical study by Molecular Dynamics and Spectroscopy": Supllementary Material

Figure SM1 - CD spectra for peptides.

Figure SM2 - Tryptophan fluorescence spectra at pH 7.4.

Figure SM3 - Dose - response curves for both peptides.

Figure SM4 - Secondary structures calculated by STRIDE for both peptides on aqueous solution.

Figure SM5 - Lifetime of H-bonds for both peptides on aqueous solution.

Figure SM6 - Secondary structures calculated by STRIDE for both peptides in the presence of the lipid bilayer.

Figure SM7 - Distance of salt briges for both peptides on aqueous solution.

Figure SM8 - Distance of salt briges for both peptides in the presence of the bilayer.

Figure SM9 - Number density profile average for both peptides.

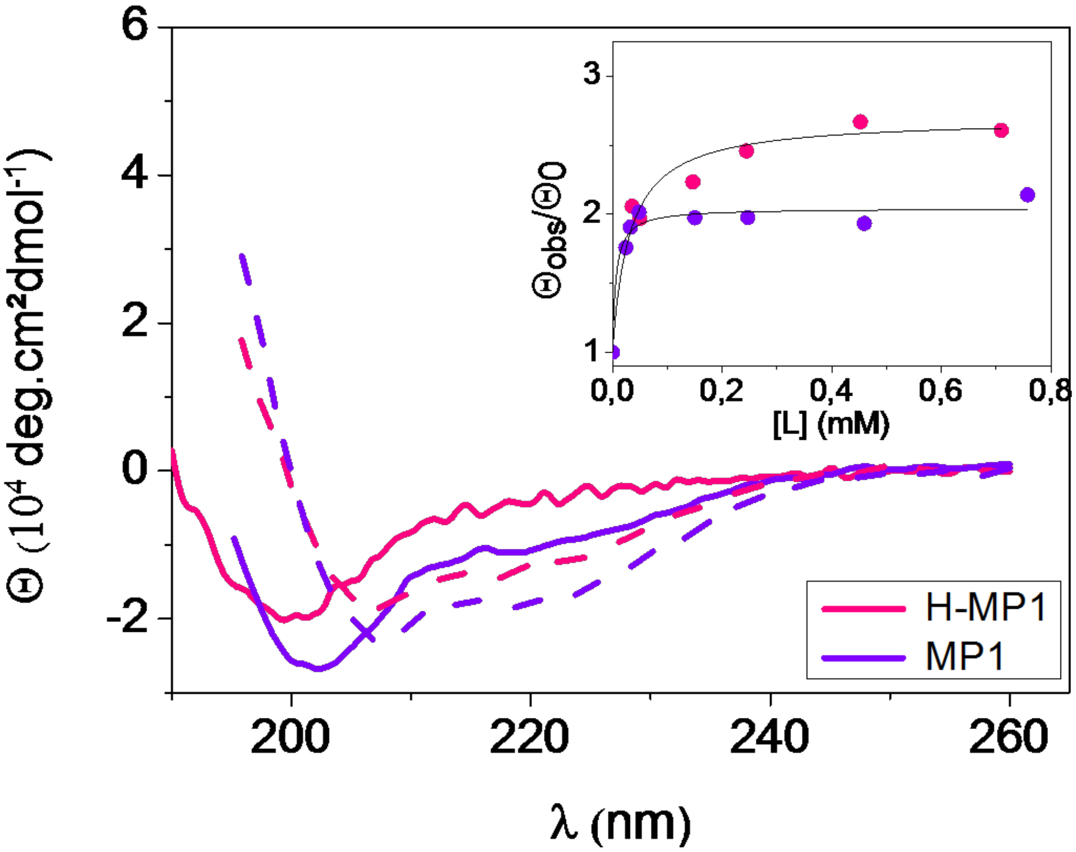


Figure SM1: (a)-CD spectra of 10$\mu$M of H-MP1 (magenta) and MP1 (purple) at pH 7.4 and 25 °C in buffer (line) and in (7POPC:3POPG) LUVs [L]/[P]=30 (trace). Inset: CD adsorption isotherms:$\theta/\theta_0$ vs total lipid concentration [L] at pH 7.4, lines are best fitting with Lagmuir model with $\gamma$=0.75

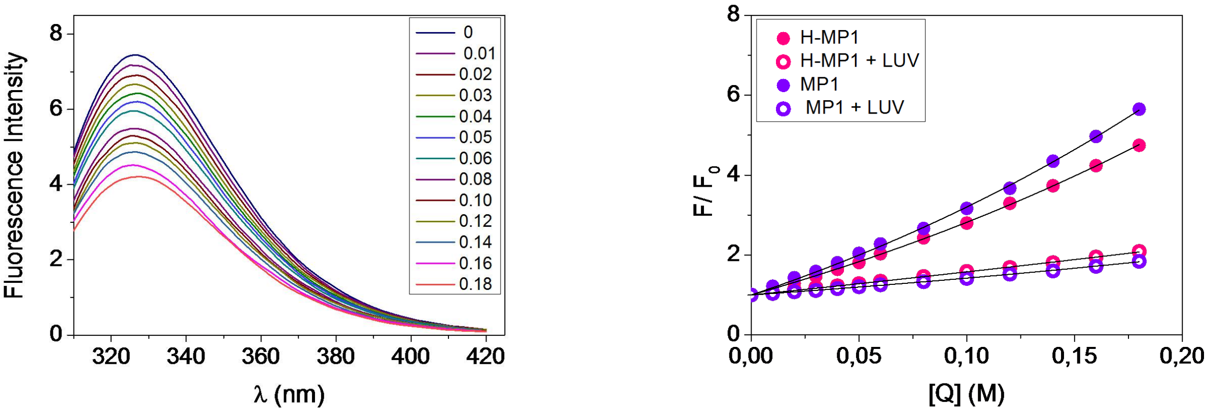


(a) (b)

Figure SM2: (a)-H-MP1 tryptophan fluorescence spectra at pH 7.4, 25 °C in the presence of acrylamide. (b) Stern-Volmer plots for the tryptophan quenching by acrylamide at pH 5.5 for H-MP1 (magenta) MP1 (purple), in buffer (closed) and in LUVs (open circles). lines are best fitting using model for collisional and static quenching (buffer) and collisional quenching (vesicles).

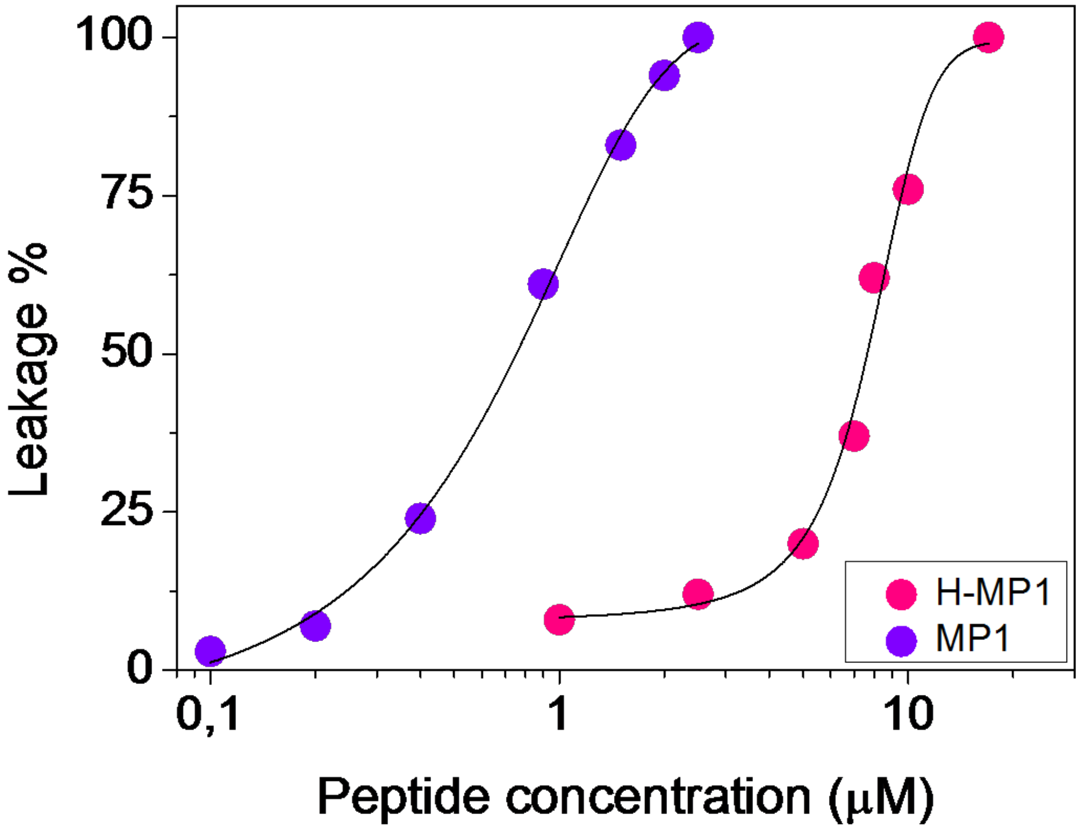


Figure SM3: Dose-response curves, % Calcein leakage from 7POPC:3POPG LUVs (100$\mu$ M lipid) induced by H-MP1 (magenta) and MP1 (purple), at pH 7.4 and 25 ° after 10 min. contact.

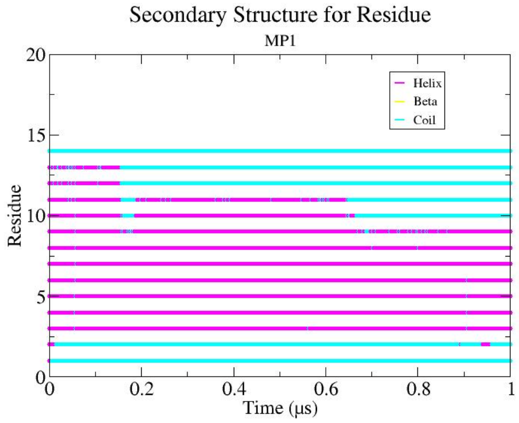


(a)

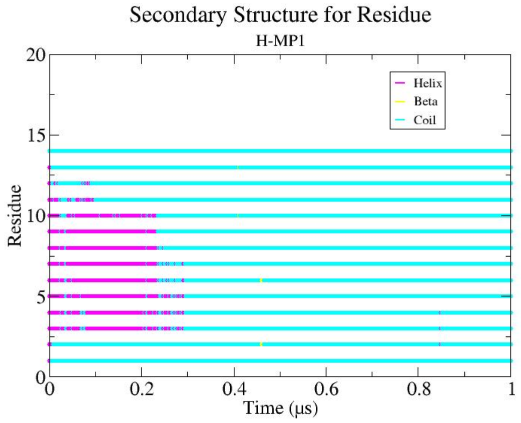


(b)

Figure SM4: Secondary structure of each amino acid during the simulation of the peptides on water, MP1 is represented on figure SM4a and HMP1 on figure SM4b, magenta for when the residue has an $\alpha$-helix secondary structure, cyan for random coil and yellow for $\beta$-sheet.

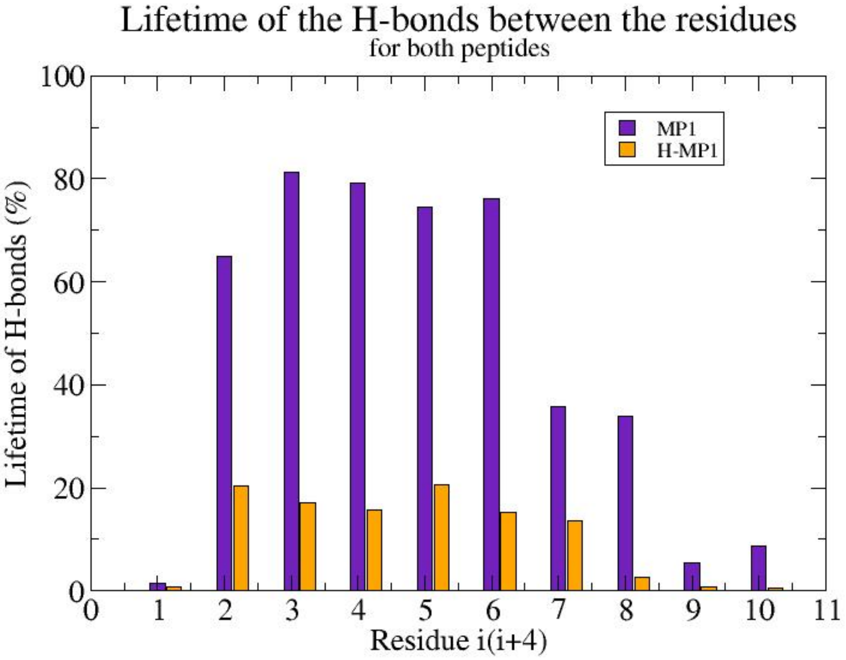


Figure SM5: Percentage of the existing hydrogen bonds between the backbone chain residues for MP1 and HMP1 on water with the purpose to see how strong is the $\alpha$-helix structuration on the peptides during the simulation. MP1 represented on purple and HMP1 on orange.

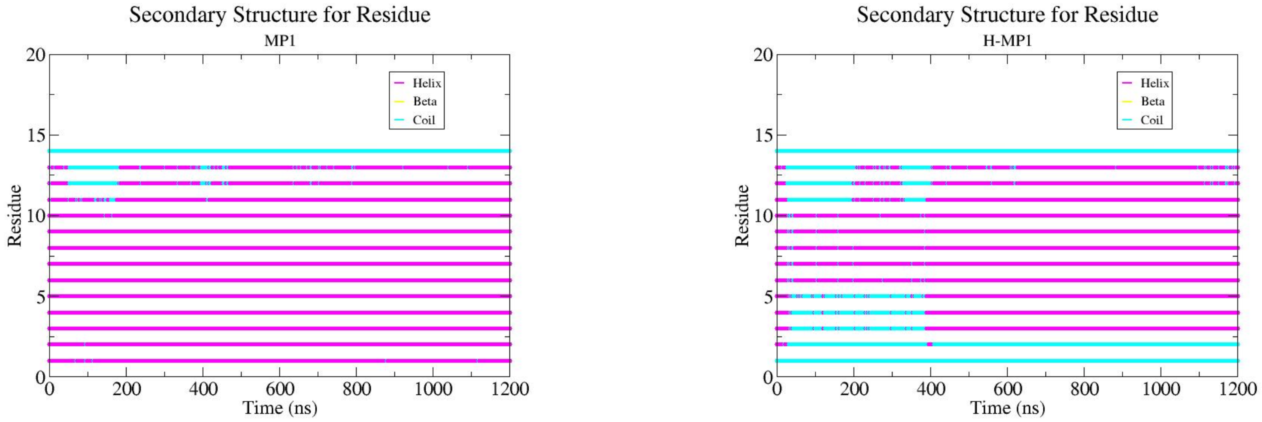


(a) (b)

Figure SM6: Secondary structure of each amino acid during the simulation, MP1 is represented on figure SM6a and HMP1 on figure SM6b, magenta for when the residue has an $\alpha$-helix secondary structure, cyan for random coil and yellow for $\beta$-sheet. The MP1 is absorbed on the bilayer starting on about 100 ns of the simulation, and the HMP1 starting on about 400 ns, its noteworthy that after the adsorption the peptide's $\alpha-$helix secondary structure increased.

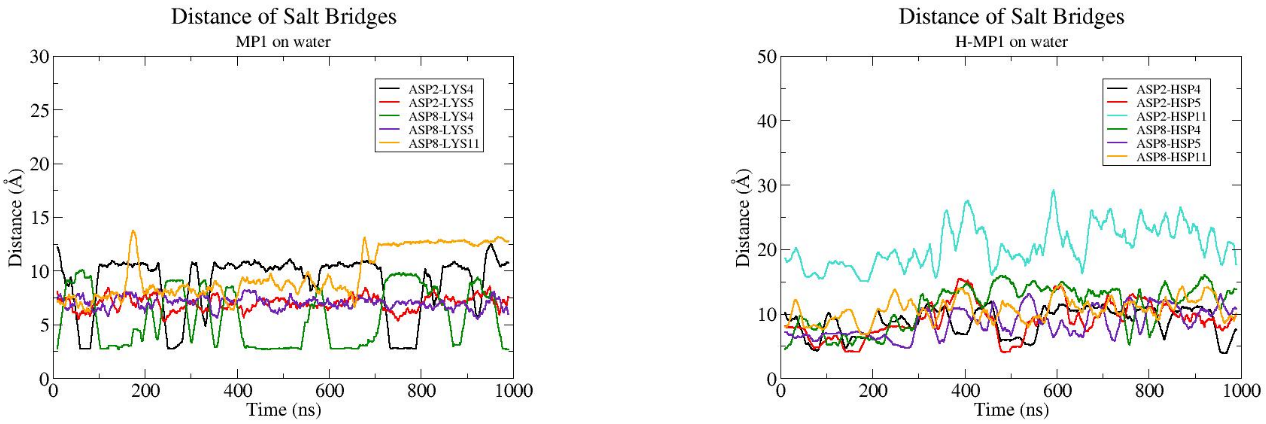


(a) (b)

Figure SM7: Distance of the salt briges between charged amino acids of peptides MP1 on figure SM7a and HMP1 on figure SM7b for simulations on aqueous solution. Bonds between residues 2 and 4 are represented on black, 2 and 5 on red, 2 and 11 on cyan, 8 and 4 on green, 8 and 5 on purple and 8 and 11 on orange.

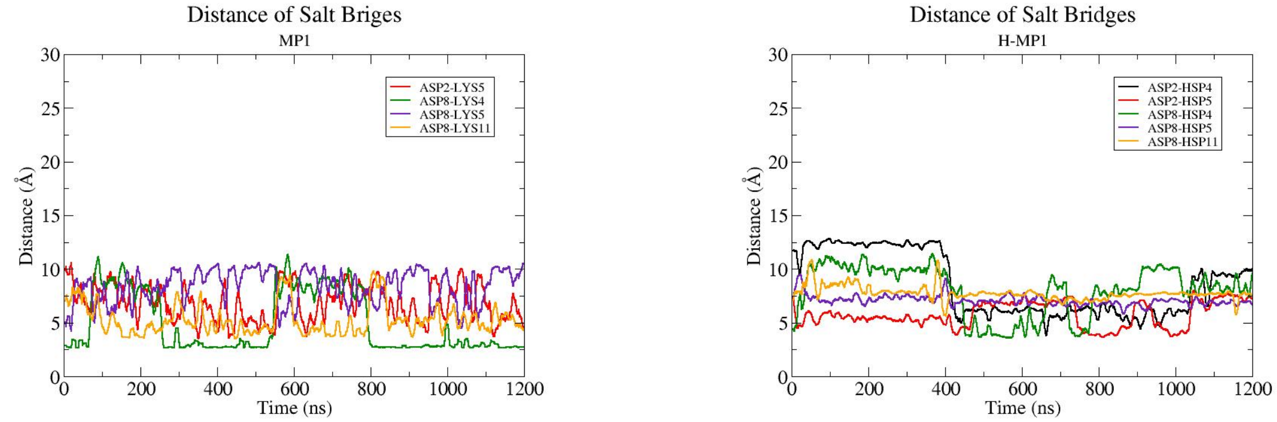


(a) (b)

Figure SM8: Distance of the salt briges between charged amino acids of peptides MP1 on figure SM8a and HMP1 on figure SM8b for the adsorption simulations. Bonds between residues 2 and 4 are represented on black, 2 and 5 on red, 8 and 4 on green, 8 and 5 on purple and 8 and 11 on orange.

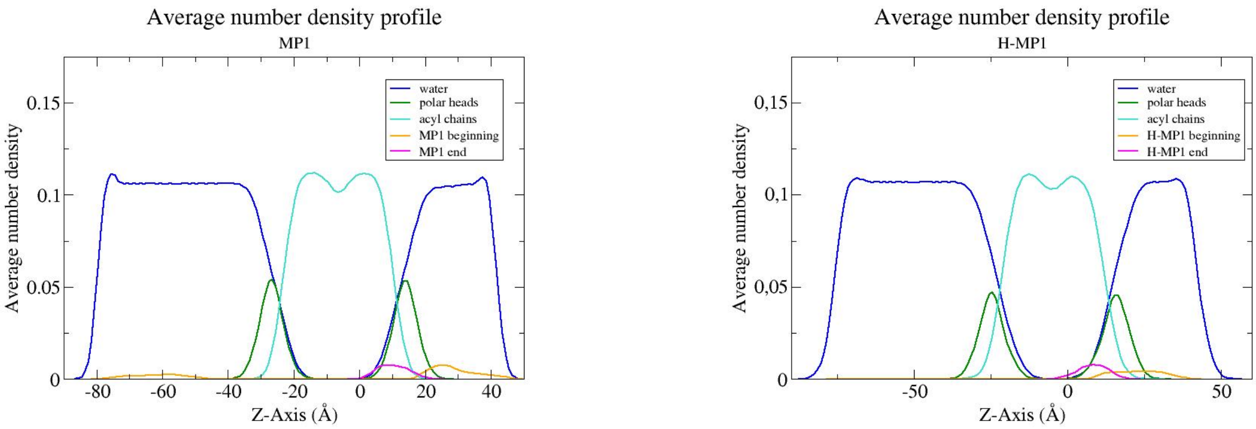


(a) , (b)

Figure SM9: Number density profile average for simulations of both peptides, MP1 on figure SM9a and HMP1 on figure SM9b, with 7POPC:3POPG bilayer in aqueous solution. Water is represented on blue, the bilayer on cyan, the peptides before the adsorption occurred on orange and after the adsorption on magenta.
